# Effects of recurrent summer droughts on arbuscular mycorrhizal and total fungal communities in experimental grasslands differing in plant diversity and community composition

**DOI:** 10.1101/2023.03.21.533628

**Authors:** Cynthia Albracht, Nico Eisenhauer, Anja Vogel, Cameron Wagg, François Buscot, Anna Heintz-Buschart

**Affiliations:** Swammerdam Institute of Life Sciences at University of Amsterdam, Amsterdam, the Netherlands; Department Soil Ecology, Helmholtz Centre for Environmental Research – UFZ, Halle (Saale), Germany; German Centre for Integrative Biodiversity Research (iDiv), Halle-Jena-Leipzig, Germany; Institute of Biology, Leipzig University, Leipzig, Germany; Department of Geography, Remote Sensing Laboratories, University of Zürich, Winterthurerstrasse 190, CH-8057, Zürich, Switzerland; Fredericton Research and Development Centre, Agriculture and Agri-Food Canada, 95 Innovation Road, Post Office Box 20280, Fredericton, E3B 4Z7, NB, Canada

**Keywords:** Biodiversity-ecosystem function relationships, experimental drought, plant-fungi interaction, AMF, Jena Experiment

## Abstract

Biodiversity loss and climate change have been determined as major global drivers affecting ecosystems and their functioning. In this context, drought was shown to have negative effects on ecosystems by disrupting ecological processes, which could be buffered in more biodiverse systems. Many studies, however, focus on effects on aboveground communities of single drought events, while dynamics of soil-borne communities are still widely unclear, despite their important roles in ecosystem functioning.

To elucidate the effect of recurrent summer drought periods on fungal communities in a long-term grassland biodiversity experiment, roof shelters were installed on grassland plots ranging in plant species richness from 1 to 16 species and plant functional group richness (1-4 groups) and composition. After 9 years of summer droughts, bulk soil was sampled and used for Illumina sequencing of the ITS2 and SSU genes to characterize the total fungal and arbuscular mycorrhizal fungal (AMF) communities, respectively.

We found shifts of AMF and total fungi community structures caused by recurrent drought and plant species richness, but no buffering of drought effects by plant diversity. Alpha-diversity (VT or ASV richness) of both AMF and total fungi increased with plant species richness but was not significantly affected by drought. Even though drought overall had minimal long-lasting effects, we found *Diversispora* and *Paraglomus* among the AMF and *Penicillium* among total fungal communities to be more abundant after the drought treatment. AMF communities were affected by the presence of individual plant functional groups, reacting stronger to presence of legumes under drought, while total fungal interaction with plant communities were similar under drought as control. AMF α-diversity differed between plant functional groups in control conditions but was independent of plant community composition under drought. In contrast, total fungi α-diversity was increased by presence of herbs and legumes only under drought.

From our results, we conclude that recurring moderate summer droughts do not strongly affect soil fungal communities. All shifts can be explained by indirect effects through the plant community and its top-down effect on soils altered by drought. Further, AMF are not less affected than total fungal communities, but rather respond differently by interacting more strongly with legumes in response to drought. Consequently, not plant species richness, but plant functional composition, dominates in shaping fungal communities under recurrent droughts.

## 1 Introduction

Plant diversity has been linked to increased productivity of ecosystems (1), but also to their stability in the face of climate extremes (2-5). For instance, in grasslands that are hot spots of biodiversity, rich in specialized plant, animal and microbe species, drought causes loss of both productivity and of biodiversity (6). Drought manipulation has been widely used to test the diversity-dependent response of grassland ecosystems’ productivity and functioning (3, 4). While drought sensitivity, subsequent loss in productivity, and recovery are dependent on precipitation patterns (7), more diverse ecosystems may be more resistant to (3, 8) and recover faster after drought (9). The diversity-stability relationship has been connected to belowground processes through quality and quantity of root exudates, litter, and root traits (10, 11). Consequently, plant diversity has been shown to positively affect soil microorganisms (12, 13) even under unfavourable conditions (14, 15).

The establishment of symbioses between plants and arbuscular mycorrhizal fungi (AMF) is considered to be one of the underlying mechanisms of the increased productivity and stability in diverse plant communities (16, 17). Mutualistic fungal communities in grasslands are dominated by arbuscular mycorrhiza, as most plant species depend on AMF for their health and growth (18, 19). The effect of drought on AMF is of special interest, because of their role in plant productivity: AMF have been shown to enhance plants’ resistance to drought across a wide range of plant species and environmental conditions (20-24). Despite their positive effects on plants, AMF themselves might be affected by global change factors, e.g. altered mycelial growth in warmed drier soils (25). In general, low soil moisture reduces microbial respiration and biomass (26). On the other hand, drought triggers an enhanced root exudation and thereby indirectly an increase of soil microbial respiration (27).

Overall, low soil moisture was found to be an even stronger factor controlling soil carbon dynamics than elevated CO2 and temperature (28, 29), although nutrient fluxes in fungi seem to be less affected by drought than in bacteria (30).,.

Interactions between AMF and plants are also affected by drought: Temperature-related alterations in spore germination and hyphal growth of AMF (25, 31) might reduce root colonisation and consequently negatively impact plant communities in the future, as root colonising dynamics change (32, 33). Even though it has been shown that root exudation is maintained during drought (34), plant-fungal interactions may only recover slowly afterwards (35). For long-term effects of drought events, de Vries, Griffiths (36) hypothesize that despite recovery of plant productivity, plant-microbe interactions remain altered through altered root exudation profiles, which also modify microbial assembly and trigger microbial activity. In grasslands, AMF identity and species richness affects plant productivity with a positive correlation between AMF and plant species richness (37, 38). Vice versa, plant identity and species richness shape AMF communities. AMF species richness can increase with plant species richness (38) and AMF community composition is shaped by plant responses to the environment (39). The community composition of AMF further shifts throughout a growing season in response to plant growth, fluctuations of root exudation and other environmental factors (40, 41). Such short-term AMF diversity studies have shown that AMF communities and plants contribute equally to shifts in their interactions switching between their multiple potential partners (42, 43). Hence, the soil-fungi-plant system should be looked at as a mutually shifting entity, responding to environmental and seasonal changes (44), including drought, to understand mechanisms of resistance and resilience of such systems. Most studies on effects of drought in grasslands, however, focus on aboveground effects or only consider general belowground proxies such as microbial biomass or microbial respiration (4, 6, 15, 45-47). More specifically, it is debated whether the diversity-stability relationship forwards a buffering effect towards belowground communities against drought. Findings vary from no buffering to increased plant diversity enhancing below-ground productivity under drought (48-50).

Ecosystem functioning relationships do not solely rely on plant species richness, but also plant functional identities as drivers of diversity effects (51, 52). Considering functional traits of plants may help with a more mechanistic understanding of BEF relationships. Distinct traits relate to functions e.g. rooting depth to acquisition of soil water and nutrients (53), and these traits are therefore determinants of diversity-productivity relationships creating a trait-axis from legumes to grasses (53). This translates to belowground communities where plant species identity and especially root traits determine the AMF community composition (54). For example, AMF respond to e.g. root diameter and consequently interact more with legumes than grasses on the root-trait axis (55) leading to legumes-associated AMF communities being distinct from those associated with grasses and herbs (56).

Our study focuses on the long-term effects of drought simulated by reducing summer precipitation to generate recurrent drought periods repeatedly for 9 years. This precipitation modification was nested in the Jena Experiment, which manipulates both plant species richness and functional group composition in a grassland ecosystem. Our investigations put a special emphasis on AMF and their interaction with plant diversity. Earlier research within the same setting showed that in the short-term, the positive relations between plant species richness and plant aboveground productivity (46) as well as soil microbial biomass (50) are maintained under drought conditions. However, this maintained higher productivity of species-rich plant communities relies on increased spring productivity to recover from growth reduction during the summer drought of the preceding year (8). As introduced above, long-term effects of prolonged drought may differ in above-ground vs. below-ground systems. While plant diversity promoted stability in aboveground productivity, it could not directly buffer summer drought effects on belowground processes such as reduced litter loss rate and microbial basal respiration and biomass, which occurred independently of plant diversity (50). We therefore aimed to expand previous findings by uncovering lasting effects of drought through analysing fungal and plant interactions one year after the last drought treatment.

Combining the potential effects of recurrent summer drought and plant diversity and functional composition on AMF and total fungal communities together, we tested the following hypotheses: i) different AMF are recruited by plants under drought, reflected also by a shift in AMF community composition and increase of AMF alpha-diversity in soil under repeated drought ; ii) importantly, because plant species richness has been shown to be an important factor for ecosystem resistance and resilience under drought, more diverse plant communities should better buffer effects of drought on belowground community compositions; alternatively, the presence or richness of plant functional groups alters drought effects; iii) AMF community responses to drought are weaker than those of other fungal taxa, and this may be due to stronger effects of plant specific shaping of AMF communities, which can be due to interactions in water or nutrient uptake.

## 2. Material and Methods

### 2.1 Experimental setup

This study was performed at the Jena Experiment field site (51) on an *Eutric Fluvisol* (FAO-Unesco 1997) developed from up to 2 m thick loamy fluvial sediments alongside the Saale river in Jena, Thuringia, Germany (50°55’N, 11°35’E, 130 a.s.l.). The mean annual precipitation is 610 mm and mean annual temperature 9.9 °C (57). The site was used as arable field for 40 years, before a conversion to grassland in 1960s, and was highly fertilized prior to the experiment setup (sowing of grassland communities) in 2002. The field site is split into 80 large plots of 20 × 20 m containing different plant community compositions varying in both species richness (1, 2, 4, 8, 16 and 60 plant species) and number of functional groups (1-4 groups: grasses, small herbs, tall herbs - summarized here as ‘herbs’ - and legumes) as described in Roscher, Schumacher (51). The plots are distributed among four blocks to account for spatial edaphic variations (including soil texture and water holding capacity) which is related to the distance of the plots to the river. Each block contains the same number of plots at each plant species richness level, covering the range of functional groups (51). Overall, there are four replicates per species richness x functional group richness combination, with plant species in different communities chosen randomly out of a pool of 60 mesophilic grassland species. The sown grassland communities are maintained by three weeding campaigns per year in March, June, and October. There are two mowing events per year, in June and September.

The drought experiment was established in 2008 as a sub-experiment nested in the pre-existing plots. Prior to the second mowing in September, transparent rainout shelters (wood and PVC sheets, 2.6 × 3 m) were installed for 6 weeks every year to induce a prolonged summer drought period (Vogel et al., 2013a) over a span of 9 years (2008-2016). Of the two sheltered subplots – hereafter referred to as treatments, one received no water after installation (‘drought’) and one received collected rain water after precipitation events (‘control’), thereby controlling for non-drought roofing effects such as altered light and temperature (58). The roof shelters excluded 39.5 mm precipitation in 2009 and reduced summer precipitation by an average of 42 % in 2008-2014 (8, 46). In July 2013, the field site was completely flooded during a natural flood event (occurring once in 200-years), and the drought treatments were continued thereafter.

### 2.2 Sampling

To gain information on the lasting effect of 8-year history of drought vs non-drought on fungal communities, bulk soil samples were taken one year after the last summer drought simulation in August 2017 from all plots except the 60-species mixes. Around 50 g of bulk soil were taken by pooling 3-5 soil cores of 0-15 cm depths per subplot. The soil was sieved at 2 mm for homogenization and to remove any plant material. Soil samples were stored at -80°C until processing for next generation sequencing. As bulk soil was sampled one year after the last summer drought treatment, two 1 g subsamples were used for measuring gravimetrically the soil water content to ensure extant drought effect. For this, 2x 1g soil were dried at 104°C with *Mettler Toledo* and *Kern DBS* moisture analyzers.

To determine drought legacy effects on plant communities, plant aboveground biomass was determined by harvesting a 0.1 m2 subplot at the end of August/begin of September 2017 by cutting plants with scissors at 3 cm above soil surface. The material was then sorted into target species, non-target weed, unidentifiable, and dead plant material and dried at 70°C for at least 48 h and weighed at weight constancy. This same procedure was also carried out throughout the 9 consecutive years of drought treatment, sampling aboveground biomass every year in May and August. Changes in aboveground plant biomass over the years were tested with ANOVA model:

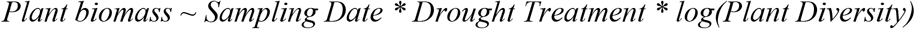

### 2.3 Library preparation and sequencing

Genomic DNA was extracted from bulk soil samples using *DNeasy PowerSoil Kit* (*Qiagen*). DNA quantity and purity was assessed using a *NanoDrop 8000* spectrophotometer (*Thermo Fisher Scientific*). The SSU rRNA region of AMF was amplified in triplicates with a nested PCR using primer pairs *Glomer 1536* and *WT0* (AATARTTGCAATGCTCTATCCCA / CGAGDWTCATTCAAATTTCTGCCC, Wubet, Weiβ (59), Morgan and Egerton-Warburton (60)) for the first and *NS31* and *AML2* (TTGGAGGGCAAGTCTGGTGCC / GAACCCAAACACTTTGGTTTCC, Simon, Lalonde (61), Lee, Lee (62)) for the second PCR step following the protocol of Wahdan, Reitz (63). To amplify the fungal ITS2 region, DNA was diluted to concentrations of 20 ng/µl and amplified in triplicates with primer pair *ITS4* (TCCTCCGCTTATTGATATGC, Ihrmark, Bodeker (64)) and *fITS7* (GTGARTCATCGAATCTTTG, Gardes and Bruns (65)) as described in (66).

Illumina sequencing libraries were prepared by purifying amplicons, barcoding and quality checking as described in Wahdan, Reitz (63) and Prada-Salcedo, Goldmann (66), respectively. Paired-end sequencing of 2 × 300 bp was performed at the *Illumina MiSeq* platform at the Department of Soil Ecology at Helmholtz Centre for Environmental Research – UFZ, Halle (Saale), Germany.

### 2.4 Bioinformatics

Raw SSU and ITS2 sequence data were filtered, trimmed, aggregated to amplicon sequence variants (ASV), and annotated with help of the *dadasnake* pipeline (67), which is based on *DADA2* (68) using amongst other softwares *cutadapt* (69) and *mothur* (70). Within the *dadasnake* pipeline, AMF sequences were classified against *SILVA 138* (71) and ITS2 sequences were classified against the *UNITE* (version 8.2) database (72) using *mothur’s* implementation of the naïve Bayes taxonomic classifier (70). The final ASV tables for target species *Glomeromycota* and *Fungi*, respectively. This resulted in a total of 2,422,141 reads with on average 16,040 (SD +-6,642) per sample of AMF. And for total fungi in 7,000,330 reads in total with an average of 46,360 (SD +-6,202) per sample.

The taxonomic annotation of the AMF ASV was not used further. Instead, AMF were clustered to virtual taxa (VTX) via *MaarjAM* (73, 74), and the read counts of SSU-ASVs assigned to the same VTX were summed up, condensing 4,865 ASVs to 114 VTX. Only SSU-ASVs that could not be assigned to a virtual taxon were extracted to construct a maximum likelihood phylogenetic tree based on a general time-reversible, discrete gamma (GTR+G) model using *MAFFT* (75) and *raxML* (76).

Thus, following the methodology of the *MaarjAM* database (73), these ASVs were assigned custom virtual taxa (VTC) with cophenetic distances below 0.03. Their read counts were merged resulting in a total of 211 virtual taxa (VTX plus VTC).

Obtained VT / ASV abundance and taxonomy results were further analysed in *R version 4*.*0*.*4* (77). Abundances were sum-normalised and VTs/ASVs that were only present in up to 3 samples and with only one read were removed.

### 2.5 Statistical analysis of fungal communities

Data was rarefied to the lowest number of reads per sample (7,350 for AMF / 27,000 for ITS) to calculate observed VT / ASV richness with *phyloseq v 1*.*34*.*0* (78). Relative abundances were generated with *transform_sample_counts* calculating sample-wise fractional abundances in *phyloseq. Jensen-Shannon Divergence* (JSD) which is effective at capturing compositional changes and commonly used in microbial ecology (79, 80), was produced with *vegan v 2*.*5-7* (81) and visualized by NMDS. Environmental factors were fitted onto the ordination with *envfit* and factors with significant correlations plotted as vectors onto the ordinations. *Vegan* was also used for PERMANOVA models (*adonis2*) and analysis of multivariate group dispersion (*betadisper*). Graphics were created with *ggplot2 v 3*.*3*.*3* (82).

We used multiple models to test for different interactions of fungal communities with their environment. Multivariate effects of experimental factors on JSD matrices were assessed with the following three models:

1.To analyse whether the experimental factors drought and plant diversity have an interactive effect on fungal communities (stratified by plot):

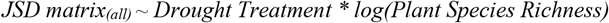

2.To test whether natural variation of soil water content across the field site adds to the drought effects (stratified by plot):

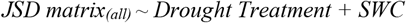

and 3. to gain a better understanding of the differences in plant-fungal-interactions between drought and control conditions, a model of effects of plant community composition within the drought / control treatment plots was tested for each treatment separately (stratified by block):

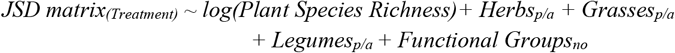

Similar models were used for *ANOVA* for α-diversity, soil moisture and plant biomass, with the exception of plot / block being added as random terms.

We calculated matrix dissimilarities of Drought and Roofed Control VT/ASV abundances from similarity matrices (1 – JSD matrix) based on the modified RV coefficient using package *iTOP* (83, 84). We analysed differential abundances of ASV/VT in drought vs control subplots with the function *ANCOM* (85).

## 3. Results

### 3.1 Long-term effects of the drought treatment on soil moisture and plant biomass

To test whether the impact of the repeated summer drought treatment was extant in the drought manipulated plots, we analysed soil water content of the bulk soil samples used for DNA extractions. Soil water content was significantly lower (t = -3.16, p < 0.05) in drought-treatment plots than in roofed controls. Soil water content was affected by the repeated drought treatments, but also by the plot position on the field site and the thereby existing variation in soil properties (Fig S1). Separately analyzing drought from control plots, we found significant effects of plant species richness and block on soil water content in both drought and control plots (Table S2). The effect size of these diversity and spatial gradients was reduced under drought.

Aboveground biomass of target plant species was determined for 2008-2016 every year in May and August. Tested against the sampling date, drought treatment and plant diversity we found plant biomass differed with sampling date (F = 17.61, p < 0.001) and plant diversity (F = 373.94, p > 0.001). However, testing biomass data of May and August separately erased the significance of the sampling date, while plant diversity effects remained, showing that the sampling date caused seasonal fluctuations instead of changes over the years. In May, plant biomass over the years was additionally affected by the interaction of drought treatment and plant diversity (F = 4.23, p = 0.04). Plant biomass significantly increased from August 2016 (last drought treatment) to August 2017 (F = 155.95, p < 0.001) influenced by plant diversity (F = 40.79, p < 0.001). This increase in biomass after the last drought treatment was consistent for all plant functional groups: legumes (F = 8.59, p > 0.01), grasses (F = 10.27, p < 0.01) and herbs (F = 10.57, p < 0.01), however, biomass of the plant functional groups individually was not affected by the drought treatment or plant diversity (Fig S5-D).

At the time of soil sampling in August 2017, total plant biomass per plot was slightly lower in former drought-exposed than in control plots, but not significantly (t = -0.64, p = 0.52). The *ANOVA* model for total plant biomass showed no effect of the drought treatment (F = 0.41, p = 0.53), but instead of the specific soil water content per plot (F = 20.72, p < 0.001; Table S2). Considering plant community composition as a factor for plant biomass, we found plant species richness to positively affect total plant biomass in both drought and control (F = 8.19, p < 0.01), with the effect size increasing under drought (F = 19.41, p < 0.001). Along the plant diversity gradient, drought did not significantly change plant biomass (neither in species-poor or species-rich communities). The absence of legumes significantly lowered plant biomass, overall (t = -4.93, p < 0.001), and the effect size of legumes on total plant biomass increased under drought (drought: F = 6.12, p = 0.02, control; F = 5.98, p = 0.02). The biomass of legumes also slightly increased under drought, as opposed to grasses and herbs that declined in biomass. The differences in plant biomass were however not significant (Table S1).

### 3.2 Community compositions

To test whether the drought and plant species and functional richness treatments have lasting effects on the composition of total fungal and AMF communities in the grassland plots, we calculated distance matrices based on Jensen-Shannon dissimilarity and conducted PERMANOVAs. The first model for drought and plant diversity interaction showed significant effects with low effect size for both the drought treatment (R2 = 0.01, p < 0.001, Fig 1-C) and plant diversity (R2 = 0.07, p < 0.001) on AMF communities and no interaction of drought and plant diversity (R2 = 0.01, p = 0.06). The variation of soil water content within the drought treatment had no significant effect on AMF communities (For the AMF community composition, the overall model for effects of drought revealed evidence of a small treatment effect (R2 = 0.01, p < 0.001) and no significance of soil water content (R2 = 0.06, p = 0.09, Fig 1-E). To assess correlation of abundance matrices of drought and control treatments, we calculated the correlation coefficient for similarity matrices (modified RV). AMF drought and control matrices showed a significant correlation coefficient (RV = 0.43, p < 0.001). Within the drought treatment plots, AMF community composition was affected by plant species richness and presence of grasses and legumes, which explained a total of 17 % of variation in the beta-diversity (Figure 1-G). Presence of herbs and number of functional groups present were not significant determining factors of AMF communities under drought. In control plots however, all plant composition factors except presence of herbs played a role in the AMF community structuring: plant species richness, presence of grasses and legumes plus the number of functional groups present amounted to 21 % variation. The low effect size of all these factors is apparent in the non-metric multidimensional scaling of the JSD matrix, as neither treatment nor plant community cause strict clustering of communities (Fig 1-A). A test for homogeneity of dispersion within AMF communities was not significant for treatments (F = 1.29, p = 0.25) nor for plant diversity (F = 2.27, p = 0.06) suggesting homogeneous dispersion of groups.

**Figure 1.**
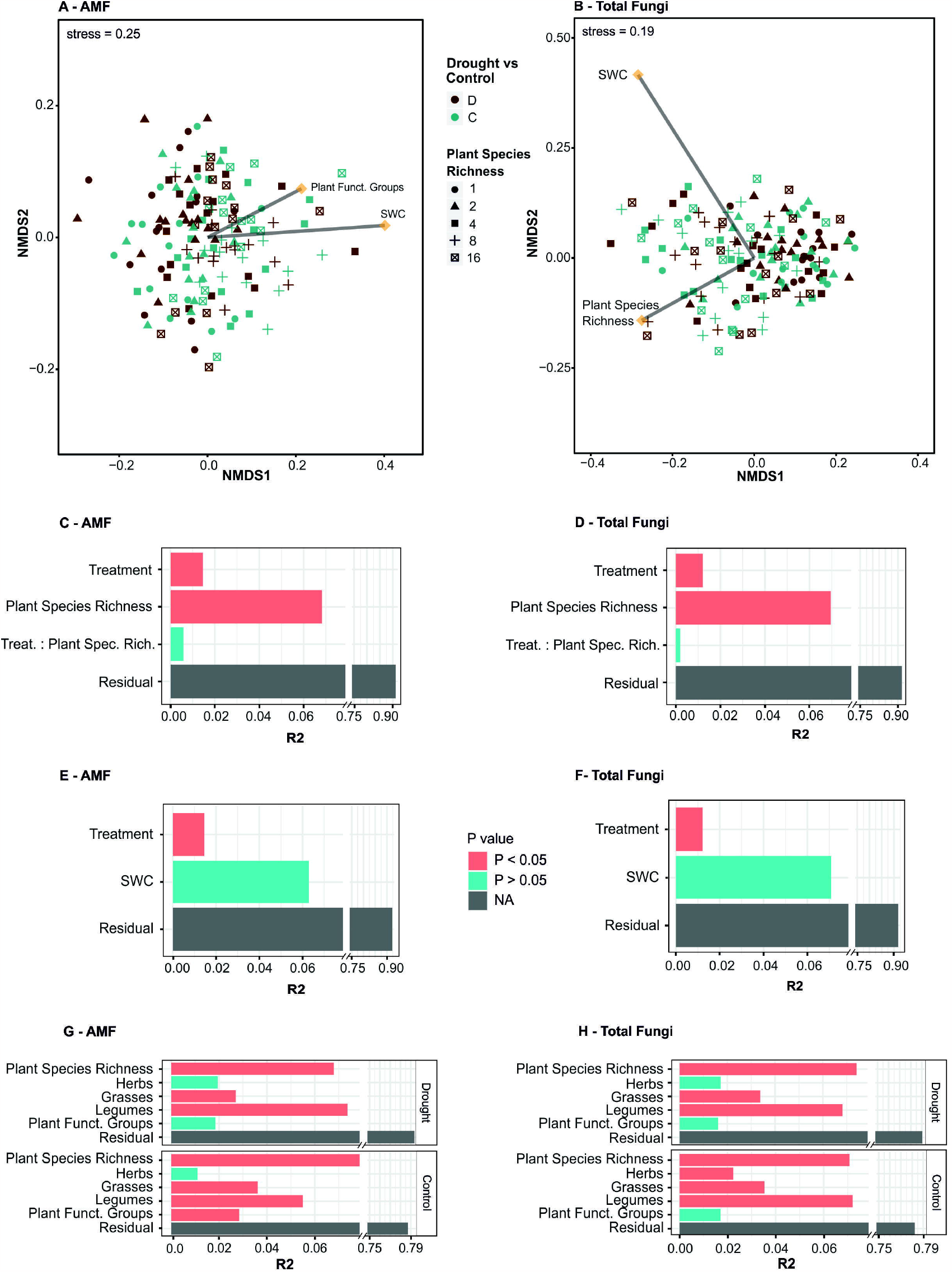
Effects of drought treatment, soil water content, and plant community composition on AMF and total fungal community composition. Non-metric multidimensional scaling of Jensen-Shannon dissimilarities for (A) AMF and (B) total fungal communities. Drought treatment is displayed by color (drought – brown, control – teal) and plant species richness by shape (1 – circle, 2 – triangle, 4-square, 8 – cross and 16 species – crossed square). Environmental factors fitted with envfit are indicated with arrows. Distance matrices were tested with PERMANOVA against effects of drought treatment and SWC (C, D) and plant community composition (E, F). PERMANOVA results are displayed as R2 value and p value (red – significant, blue – non-significant).

For the beta-diversity of the total fungal communities (Fig 1-B), the first model on drought and plant diversity interactions revealed significant effects with low effects size for the drought treatment (R2 = 0.01, p < 0.001, Fig 1-D) and plant diversity (R2 = 0.07, p < 0.001), however these two factors do not interact (R2 = 0.00, p = 0.63). The variation in soil water content tested with PERMANOVA model 2 does not add additional variation to total fungal communities (R2 = 0.07, p = 0.20, Fig 1-F). The coefficient of correlation between drought and control similarity matrices was higher than in the case of AMF with RV = 0.74 (p < 0.001). Within the drought exposed plots, plant species richness, presence of grasses and legumes, but not herbs, shaped total fungal communities with a total explanatory value of 17.6 %. Total fungal communities under drought were not affected by the number of functional groups present (R2 = 0.07, p = 0.11). The influence of these plant community factors was similar in significance and effect size in control plots (Fig 1-H), however in control plots herbs also determined fungal community structures (R2 = 0.02, p = 0.02). Tests for homogeneity of dispersion within ITS communities were not significant for the drought treatment (F = 0.22, p = 0.64), but significant for plant species richness (F = 3.34, p = 0.01). An adhoc TukeyHSD further revealed different dispersion between groups of 4 and 1 plant species (padj = 0.05) and 16 and 1 plant species (padj = 0.05), suggesting that parts of the observed differences were due to differences in dispersion rather than between the treatments.

Differential abundances were analysed with ANCOM. Among the AMF were only the two taxa VTX00354 (*Diversispora Torrecillas 12b*) and VTC00015 (*Paraglomus sp*) significantly differnetialy abundant – both higher abundant under drought with log fold changes of 1.15 (VTC00015) and 1.21 (VTX00354). Within the total fungal communities, we found three taxa that were significantly higher abundant under drought conditions: *Penicillium jenensii* with a log fold change of 1.85, an unidentified *Penicillium sp*. (log fold change = 0.33) and an unidentified *Ascomycot sp*. (log fold change = 1.02).

In brief, AMF and total fungal communities were shaped by plant community composition and the drought treatment caused lasting shifts in community structures. However, drought and plant community explained only small proportions of the AMF and total fungi community structures.

### 3.3Drought and plant diversity effects on α-diversity of AMF and total fungal communities

Given the observed effect on community composition, we also tested whether the α-diversity of AMF and total fungi was affected by the repeated drought treatment. Paired t-tests of VT / ASV richness in drought and control subplots showed no significant differences for neither AMF (t = 0.39, p-value = 0.69) nor total fungal (t = 1.70, p = 0.09) communities. Testing α-diversities against model 1, AMF VT richness was neither significantly altered by drought (F = 0.13, p = 0.72) nor plant diversity (F = 0.12, p =0.73) nor their interaction (F = 2.23, p = 0.14). Total Fungi ASV richness was not altered by drought (F = 1.29, p = 0.26) or an interaction of drought and plant diversity (F = 0.02, p = 0.88), but showed marginal significance for a plant diversity effect (F = 4.08, p = 0.05).

When applying ANOVA with the drought treatment model to α-diversity, neither drought treatment as a factor nor specific SWC were influencing α-diversity of AMF or total fungi (Fig 2, Table S3). Regarding the influence of plant richness and presence of functional groups on fungal α-diversity, plant species richness (R2 = 0.13, p = 0.03) and presence of herbs (R2 = 0.05, p = 0.03) increased α-diversity of AMF in control plots, but no plant functional group was significant for AMF α-diversity under drought (Table S3). In contrast, the presence of herbs (R2 = 0.06, p = 0.03) and legumes (R2 = 0.05, p = 0.04) significantly increased total fungal communities under drought, but not in control plots. The fungal α-diversities in more diverse plant communities were not affected by the drought treatment.

**Figure 2.**
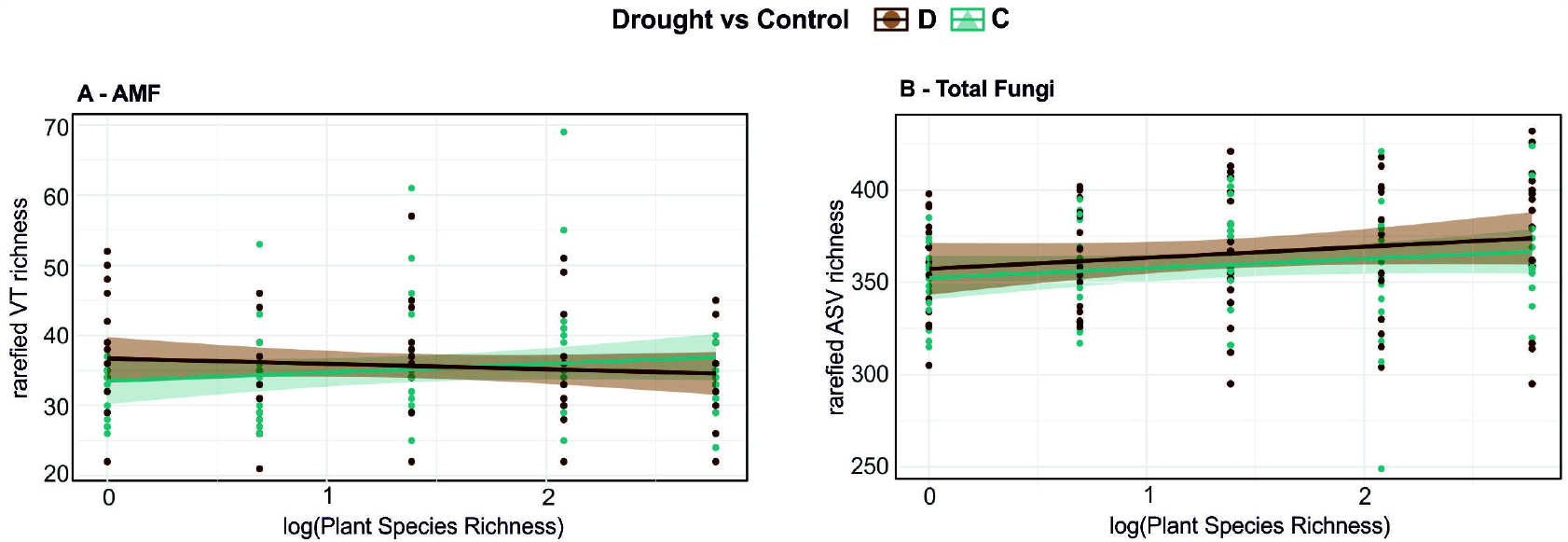
Alpha-diversity of AMF and total fungal communities according to plant species levels and treatments. (A) Observed AMF VT richness and (B) total fungal ASV richness after rarefaction versus log-transformed plant species richness (1 – 16). Treatments: brown – drought; teal – control subplots.

**Figure 3.**
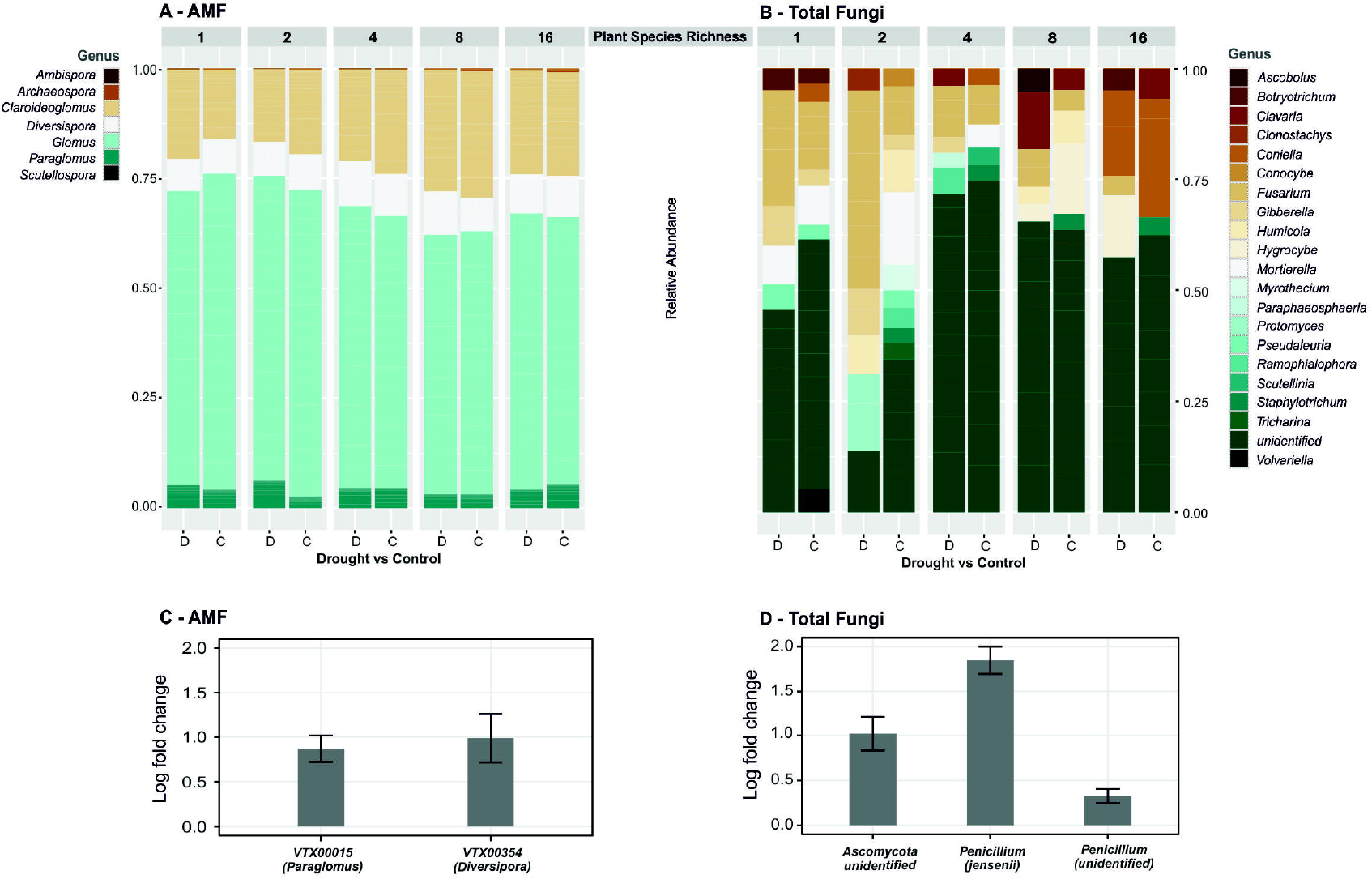
Genus level composition and changes of AMF and total fungal communities. Relative abundances of (A) VT of AMF and (B) ASV of total fungal communities > 5 %, subdivided by the plant species richness levels (1-16) in drought [D] vs control [C] plots. Changes in significantly differential abundant taxa (C – AMF, D – total fungi), presented as log fold change of abundance after drought compared to control plots with standard deviations bar.

In short, drought had no major impact on fungal α-diversity. Plant functional groups played a role under drought for total fungal communities, but not for AMF. AMF α-diversity however was affected by plant communities in control plots.

## 4.Discussion

### 4.1 Repeated summer drought has lasting effects on soil water content but not on AMF α-diversity

The 9-year history of drought treatment effectively changed the soil water content, which appeared still significantly lower in plots that were exposed to repeated drought even one year after the last treatment. Previously, Vogel, Fester (58) found soil moisture to not differ significantly between the treatments in the same experiment after one year, but found a response of soil moisture to the drought treatment over time, as the difference was significant in the third year of drought treatment. This is congruent with the soil water content being still divergent after 8 years of treatment, despite the field site flooding in 2013 (86). With an 8-year history of summer drought, the reduction amounted to only about 2 % less soil water in the drought treatment compared to control plots. SWC additionally varied across the field site, coinciding with edaphic variation of the field site (87) which was accounted for by always including the field block as error term. Overall this shows that even with a moderate, but repeated, drought treatment, changes in ecosystems become visible, and we need even more experimental approaches of realistic climate scenarios as opposed to climate extremes (88) to understand consequences on multiple ecosystems levels.

Arbuscular mycorrhiza have repeatedly been shown to enhance drought tolerance of their plant partners (89). Previous studies have shown that plants rely on mycorrhizal symbiosis under drought stress, but that AMF colonisation decreases under drought (90, 91). Soil moisture regulates AMF community assembly, with drought causing shifts in relative abundances including the absence of certain AMF, which results in a reduced α-diversity (92). Therefore, we expected to see changes in the α-diversity of AMF in response to repeated drought. Our treatment with eight years of prolonged summer droughts, however, had no significant impact on AMF α-diversity (VT richness). Neither the repeated drought treatment nor the resultant SWC had significant effects on AMF α-diversity, therefore drought in our experiment did not cause a recruitment of more species. Analyses of the root and rhizosphere mycorrhiza may yield stronger α-diversity effects because they represent the place of AMF-plant interaction. In this regard, we also have to take into account that the flooding of the field mid-experiment might have been a mode of transport for propagules.

Despite the α-diversity of AMF remaining similar under drought across the diversity treatments, the community composition did change. This effect was apparent regardless of the high variation of SWC along the field and across the plant diversity gradient. However, the variation explained by drought was limited, which is in line with AMF having been shown able to endure drought (36). Two AMF VT were found to have a higher relative abundance under drought. The first one was identified as *Diversispora*, whose mycorrhizal infection is inhibited by drought (93). But *Diversispora* were found to be present predominantly outside rather than inside plant roots (94, 95) exhibiting low infection rates of roots but proofing effective for plant productivity (96). These edaphophilic AMF show high biomass allocation to extraradical hyphae and increase plant nutrient uptake, but are usually characterized as sensitive to drought (97). With moderate drought as we had in our experiment, this drought sensitivity of *Diversispora* might however have been overruled by their interaction with plants. The second AMF genus with increased abundance under drought was *Paraglomus*. This genus is rhizophilic with high abundance in grass roots (97) and was previously reported as having a positive influence on drought resistance of plants (98, 99). A higher relative abundance of these two genera under drought suggests distinct responses of AMF species in relation to their host preference and their drought response. However, this is in terms of relative not quantitative abundance and might also be results of these genera being persistent while others decrease.

Summing up the effects of repeated summer drought treatment on AMF communities, we found minimal drought-induced changes on AMF α- and β-diversity, but the differential abundances of an edaphophilic and rhizophilic AMF indicate shifts in the interaction of AMF with plant partners. A reason for the relatively small drought effects may be the sampling of bulk instead of rhizosphere soil. Even though bulk soil in grasslands is still in vicinity of and highly shaped by plant roots (100), we cannot be sure that our results would be identical in the rhizo- or endosphere. Rhizosphere soil is subject to stronger changes through plant nutrients and biomass and other microbes like bacteria (101) and even in the confines of pot-experiments has been reported to respond stronger to drought stress than bulk soil (102). Nevertheless, resistance of bulk soil communities is important for potential feedback on plant communities because they represent the general pool of available interaction partners.

### 4.2 More diverse plant communities do not buffer effects of drought on belowground communities

Recurring drought events affect ecosystems negatively, however plant species richness proved to be an important factor of resistance and alleviation (3, 4, 8, 46, 103). As Wagg, O’Brien (8) reported, the plots with high plant diversity maintained long-term productivity under drought in our experiment.

Similarly, in the year after the last drought treatment, we found aboveground plant biomass to be lower after repeated drought with the reduction of productivity tending to be higher in less diverse plant communities. The influence of plant diversity as well as presence of legumes on biomass increased under drought. Noteworthy, legumes were the only plant functional group with an increase in biomass after drought exposure.

Throughout the repeated drought treatments, plant biomass was overall not greatly reduced by drought. Drought induced seasonal shifts: May more susceptible to treatment and treatment*diversity interaction as already shown for the first 5 years by Wagg, O’Brien (8) It seems especially higher divers plant communities had stumped productivity and could restart quick after last drought.

However, drought exposed plant communities recovered quickly after the last drought exposure. This is in line with findings of Chen, Vogel (104) who collected seeds from this same experiment, and exposed the plants of drought or ambient legacy once again to drought and found drought-legacy plants to recover much quicker after droughts than ambient-legacy plants.

By sampling a year after the last drought treatment, we tested for a drought legacy effect rather than a drought response. Plants have been shown to adapt to drought stress and even establish a drought memory with physiological and morphological alterations (105). Changes in precipitation patterns can reduce net primary production and affect e.g. soil respiration and nitrogen mineralization (106). Microbial processes respond quickly to changes in soil moisture, but can revert back just as quickly (105). Contingently, the shifts in microbial communities we found are a result of the plants’ drought memory.

With plants shaping belowground processes, we expected to see a positive effect of plant species richness on AMF community compositions in both control and drought plots and a buffering effect against the drought effect by higher plant diversity. Correlations of the distances between community compositions as well as the PERMANOVA models showed that total fungi and AMF were affected by plant species richness and community composition, independent of drought events. Likewise, our results suggest that a higher plant diversity does not buffer negative drought effects on soil fungi diversity. This lack of effect might reflect that the drought effect on soil moisture was weak along the plant diversity gradient. In addition, while soil under richer plant communities had a higher water content in control plots, this relationship decreased under drought, indicating that the long-term buffering effect of diverse plant communities on the soil water content was also weak. Finally, plant species richness was an important factor for β-diversity as well in controls as under drought stress.

Our results are in accordance with data from the beginning of the experiment (50) showing that after one year of drought, higher plant diversity did not buffer against belowground drought effects, such as lower litter decomposition rates and microbial respiration. These findings suggest that plants in high diversity settings maintain their aboveground rather than belowground productivity under short-term adverse conditions. In our study, plant biomass was not significantly reduced under drought at any plant diversity level. However, increasing effect size of plant species richness on biomass under drought indicate stronger differences between the diversity levels. The described maintenance of plant productivity could be explained by short-term variation in biomass production, which increased in more diverse communities due to compensatory recovery from drought (8). Considering this drought-induced short-term variation, buffering effects of diversity might become more visible with seasonal variations, and Wagg, O’Brien (8) found spring growth to compensate for summer drought induced productivity losses but only at high plant diversity. Our data comes from just a single time point in the driest month, which may have caused us to miss temporary changes. However, we would expect those short-term changes to occur predominantly in the rhizo- or endosphere, while the bulk soil microbiome reacts less to droughts (102, 107).

Correlations of AMF abundance and plant species richness have been previously shown, for example, AMF abundance decreasing in the presence of certain grass species and increasing in the presence of legumes (55). Legumes have been proven to facilitate carbon allocation to fungi under moderate drought stress (108). Our results showed that the number of different functional groups is only relevant for α-diversity of AMF in drought but not control plots, while it shaped β-diversity only in controls. However, presence of grasses and legumes promote AMF communities in both control and drought with the influence of legumes even increasing under drought. The reduced effect of grasses on AMF communities might stem from grasses raising the soil water content in the short term (50), an effect which did not persist to the end of the experiment according to our soil water content measurement. Kivlin, Mann (54) found plant identity to be a major factor in shaping fungal communities despite environmental gradients, but diverse climate changes can interrupt plant-microbe interactions (109). Drought in particular affects water uptake, nutrient concentration such as C:N ratio, extending to increased plant-plant competition (110). Some functional plant groups benefit more from AMF, as Chagnon, Bradley (111) suggest: limitation of carbon allocation to root exudates caused by stress negatively affects plant-soil interactions, but slow growing plants invest more in AMF and receive more long-term benefits. Additionally, AMF have varying effects on productivity of different plant functional groups. Legumes seem to benefit in terms of whole plant growth while mycorrhization for herbs increases reproductive growth (112). This may explain the stronger effect of the presence of legumes on AMF communities. Legumes use more water, but have a higher water use efficiency than e.g. grasses (113) and together with their facilitated carbon allocation towards fungi under drought become a more desirable interaction partner for fungi. Microbes vice versa intensifying plant-plant competition under drought (110) could further explain the increase in legume biomass and legume-fungal interaction confirming long-term benefits of AMF symbiosis. AMF have host preferences (43) which might increase under stress conditions in relation to available transferable nutrients. Interestingly, AMF α-diversity was higher in monocultures even though AMF host-preferences would suggest higher AMF diversity with increasing plant diversity (114, 115).

However, monocultures seem to benefit from AMF colonization even under drought, while plant mixtures might encounter disadvantages from demanding AMF species (116).

### 4.3 AMF response to drought is hardly weaker than that of other fungi

To test whether AMF as biotrophic interaction partners of plants are specific in their response to drought, we compared their response to that of other fungi. *Glomeromycota* made up 3.5 % of abundance in control plots within the ITS2 dataset and increased to 4.1 % of abundance under drought. This increase was expected: Even though fungal communities in general are rather resistant to drought (30), they have been shown to shift towards a lowered abundance in *Agaricomycetes* and other fungal groups and a higher abundance in *Glomeromycota* (117). However, the low proportion of *Glomeromycota* is due to the choice of the general fungal ITS2 primers, which bias amplification against the *Glomeromycota* (118, 119). In addition, most ITS2-reads of *Glomeromycota* could not be identified at the genus level. Therefore, the results discussed in the previous sections stem from AMF-specific amplifications and the rest of the community was assessed independently.

The α-diversity of total fungal communities did not change in response to drought, as for the AMF. But the explanatory value of legumes for general fungi increased under drought, indicating that the increasing biomass of legumes under drought plays a role for general fungal diversity and maybe a shift towards more AMF species. Sweeney, de Vries (55) showed not only that AMF have a preference for legumes driven by root traits, but also that higher abundance of AMF in roots leads to a reduced abundance of fungal pathotrophs.

Both AMF and the total fungi β-diversities did minimally change under repeated drought, but they differed in their response to plant communities under adverse conditions. In the control plots we found AMF to be more responsive to plant species richness than the total fungi communities. The effect of legumes and herbs was similar for both fungal communities, but total fungi are additionally shaped by grasses, while for AMF communities the plant functional diversity plays a role. Total fungi communities under ambient and stress conditions were consistently affected by plant diversity and functional composition. This is in line with several studies showing that the fungal community is driven by plant functional identity and root traits with e.g. saprotrophic fungi being less abundant in grass roots (55) and plant functional diversity being a major driver for fungal richness and community structures even under stress conditions like drought (120, 121).

While we found two significantly higher abundant AMF VT under drought, the total fungi community did not detect any differential AMF at genus level. Here we found *Penicillium* to be more abundant under drought and no genera significantly decreasing under drought stress. One of the two *Penicillium, P. jensenii*, is often found in locations with higher moisture (122), but as saprophytic and phosphate-solubilizing fungi they might thrive under drought as more dead plant matter becomes available (123).

Comparing the responses of AMF and total fungi in our experiment, AMF communities displayed stronger shifts under drought than the total fungal communities. The drought-induced shifts were towards a slightly higher richness and abundance of AMF. Juxtaposing AMF communities and total fungi communities emphasizes even more the importance of plant functional identity as driver of fungal soil communities and plant-fungal interactions.

### 4.4 Conclusion

In summary, the effect of eight consecutive years of prolonged summer drought events on fungal communities was minimal across the plant diversity gradient. This means soil-borne fungi, including AMF, were similarly resilient to drought at lower and higher plant species richness. Fungal α-diversity was not negatively affected by drought, and beta-diversity of AMF and total fungi communities only slightly shifted. The fungal communities therefore did not seem to be protected from drought by plant diversity, in contrast to plant productivity, which was more strongly affected by plant species richness under drought. All these findings suggest that plant diversity itself does not shape fungal community structures under drought stress. We also found that the influence of plant functional diversity on AMF communities decreased under drought, while it increased for general fungi. However, the effect of the presence of legumes increased under drought. While overall plant biomass productivity did not change greatly in response to repeated yet moderate drought exposure, shifts we do find in fungal communities indicate indirect drought effects caused by changes in plant-fungal interactions. This study illustrates the range of disturbances droughts cause on multiple levels of ecosystems, also in moderate drought scenarios.

## 6 Author Contributions

A.V., C.W. and N.E. conceived and carried out the field experiment. C.A. conducted sample preparations and statistical analyses. C.A., A.H.-B. and F.B. wrote the manuscript with input from all authors.

## 8 Acknowledgments

We thank the technical staff Steffen Eismann, Ute Köber, Gerlinde Kratzsch, Katja Kunze and Heike Scheffler for their work in maintaining the field site and also many student helpers for weeding of the experimental plots. Further, we thank the central coordination team Anne Ebeling and Annette Jesch, and the data management team. We further thank Alexandra Weigelt and Michael Scherer-Lorenzen who were involved in the design of the experiment and Anne Ebeling and Holger Schielzeth for assisting with statistics and the writing process. The sequencing data were processed at the High-Performance Computing (HPC) Cluster EVE, a joint effort of both the Helmholtz Centre for Environmental Research - UFZ and the German Centre for Integrative Biodiversity Research (iDiv) Halle-Jena-Leipzig, whose administrators we thank for excellent support.

## 9 Data Availability Statement

The datasets generated for this study can be found in the NCBI SRA under accession number PRJNA914200.

