## Supplementary Material for "Effects of recurrent summer droughts on arbuscular mycorrhizal and total fungal communities in experimental grasslands differing in plant diversity and community composition"

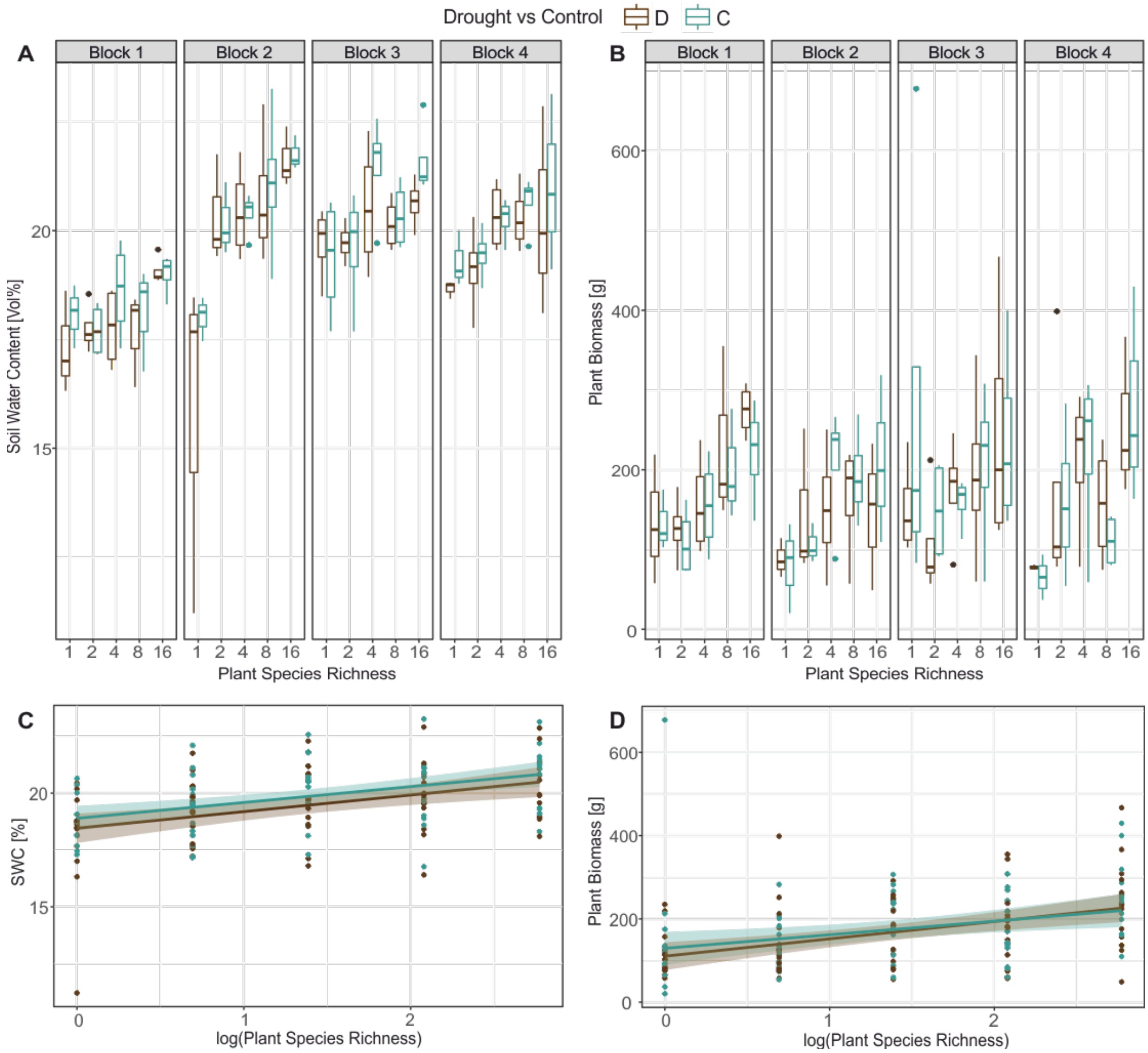


**Supplementary Figure S1.** Soil water content (SWC) and total plant biomass in control and drought exposed plots of different plant richnesses across the four field site blocks. Variation of soil water content (%) per block and plant species richness as boxplot (A) and linear model (C). Aboveground plant biomass (g) per block and plant species richness as boxplot (B) and linear model (D).

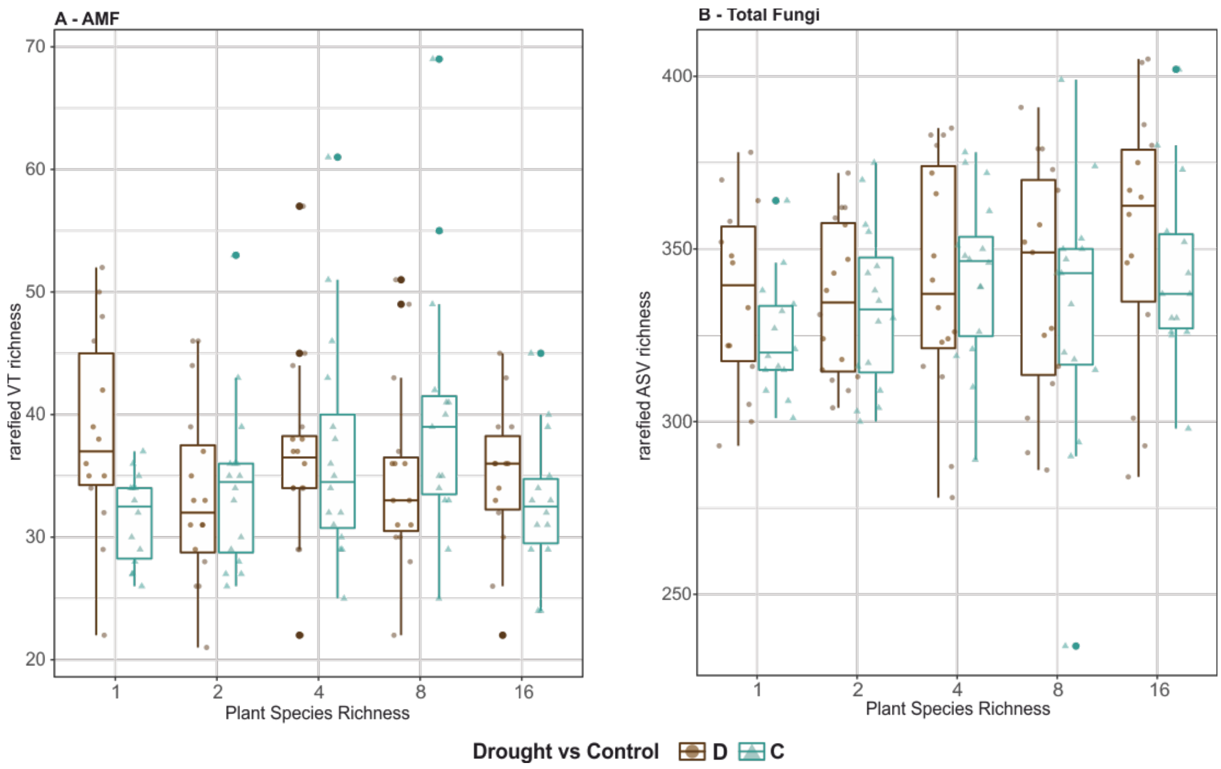


**Supplementary Figure S2.** alpha-diversity of AMF and total fungal communities according to plant species levels and treatments. (A) Observed AMF VT richness and (B) total fungal ASV richness after rarefaction, subdivided by plant species richness (1 – 16). Treatments: brown/circle – drought; teal/triangle – control subplots.

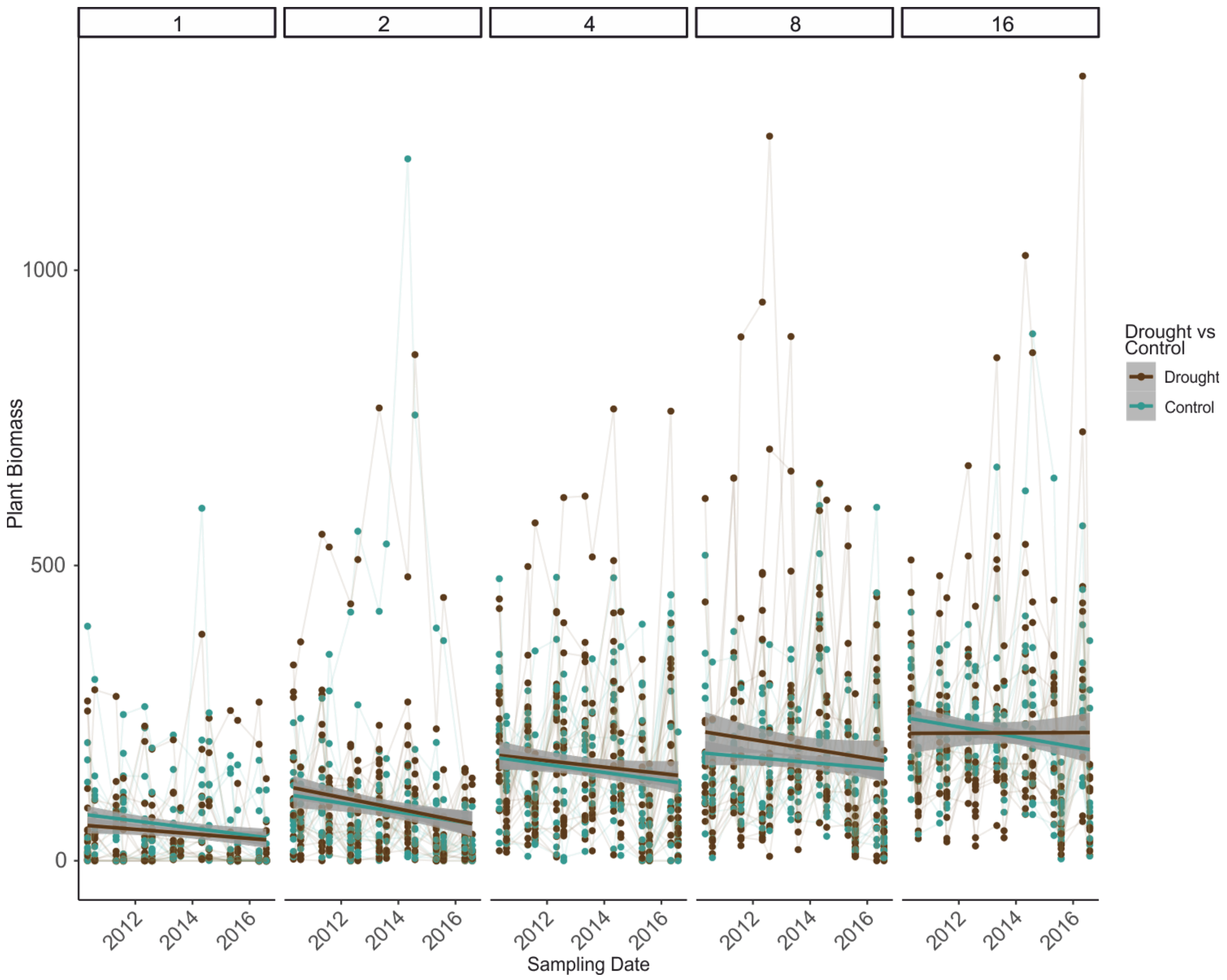


**Supplementary Figure S3.** Plant Biomass [g] of grassland communities under drought and ambient conditions over the years separated by plant diversity. Between 2010 and 2016 every May and August aboveground biomass was determined by cutting samples from a 0.1 m$^2$ subplot. The plant material was sorted into target species, weeds, and dead plant material, dried and weighed. Data points from the same plot are connected.

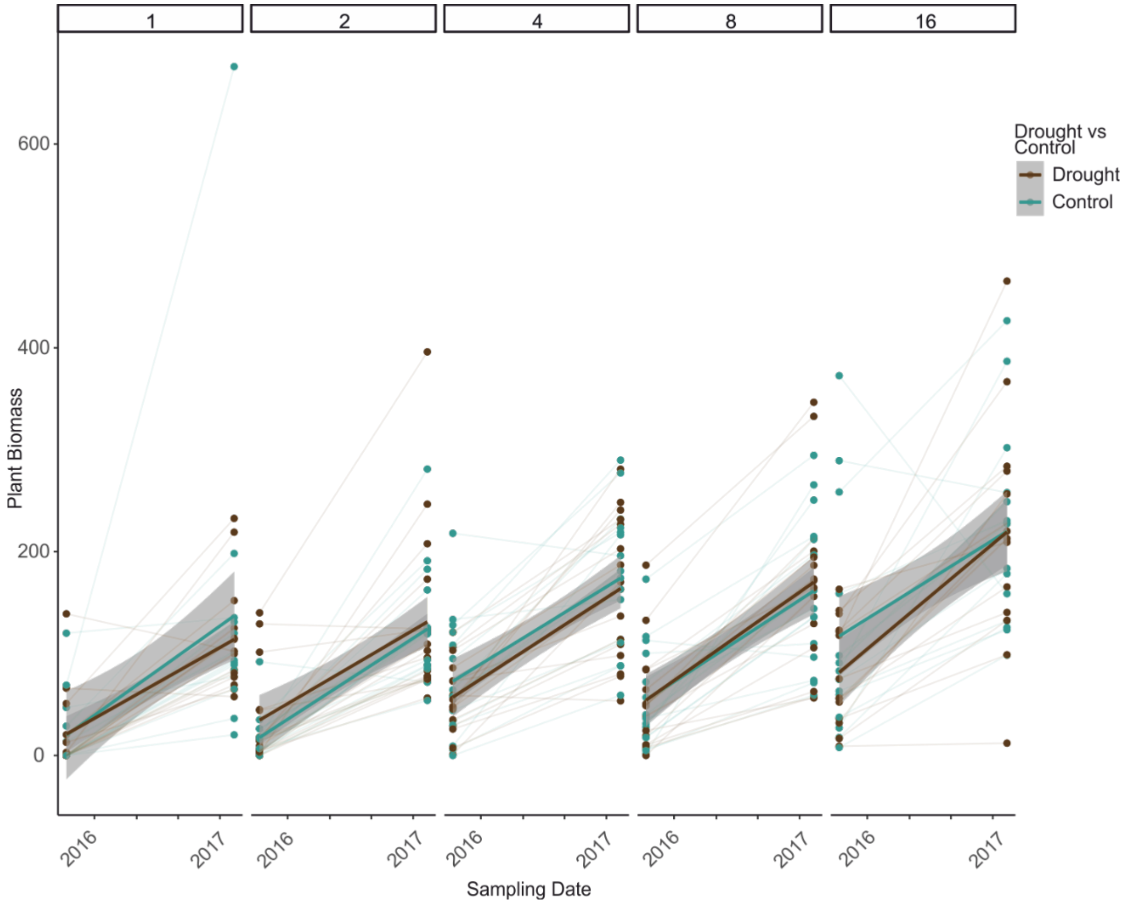


**Supplementary Figure S4.** Plant Biomass of grassland communities under drought and ambient conditions in August of the last of 8 consecutive years of drought treatments (2016) and August one year after the last treatment (2017). Aboveground biomass was determined by cutting samples from a 0.1 $m^2$ subplot. The plant material was sorted into target species, weeds, and dead plant material, dried and weighed.

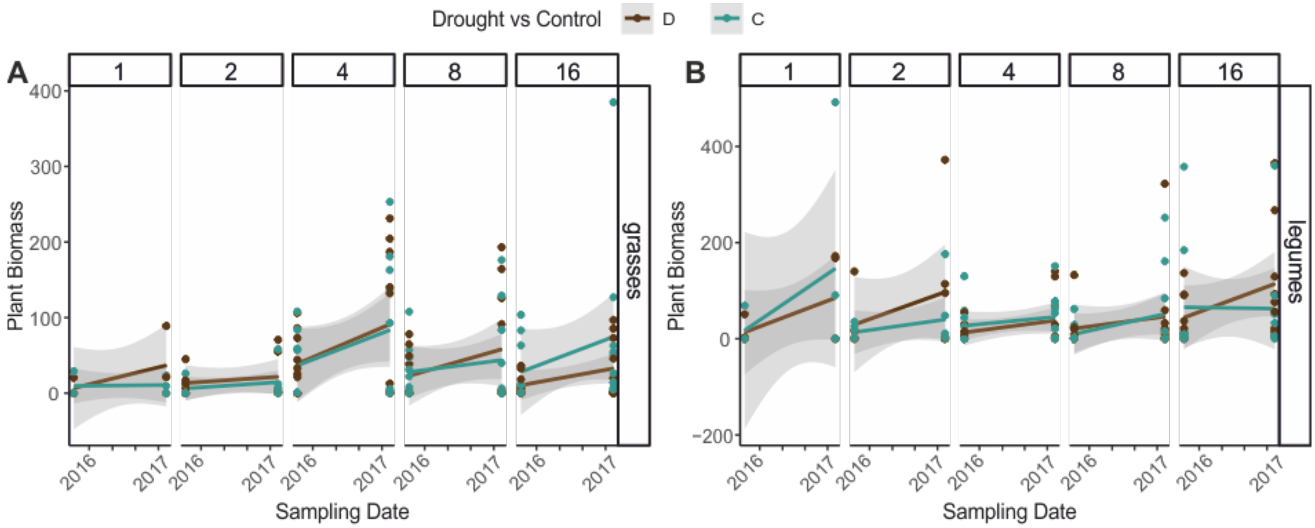


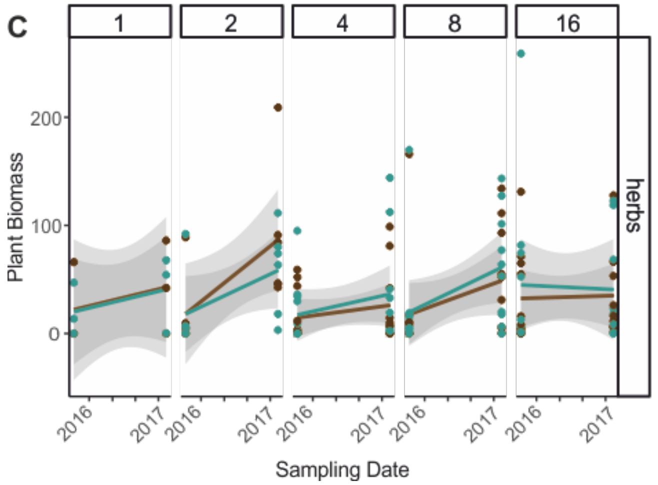


D

| | grasses | | legumes | | herb | |
|---|---|---|---|---|---|---|
| | F | Pr(>F) | F | Pr(>F) | F | Pr(>F) |
| **Date** | 10.27 | **< 0.01** | 8.59 | **< 0.01** | 10.58 | **< 0.01** |
| **Drought treatment (DR)** | 0.11 | 0.74 | 0.08 | 0.78 | 0.24 | 0.63 |
| **Plant Diversity (Div)** | 1.08 | 0.30 | 0.25 | 0.62 | 0.15 | 0.70 |
| **Date : DR** | 0.02 | 0.90 | 0.31 | 0.58 | 0.01 | 0.91 |
| **Date : Div** | 0.29 | 0.59 | 1.09 | 0.30 | 2.17 | 0.14 |
| **DR : Div** | 1.73 | 0.19 | 0.18 | 0.67 | 0.66 | 0.42 |
| **Date : DR : Div** | 0.31 | 0.58 | 0.75 | 0.39 | 0.06 | 0.81 |

**Supplementary Figure S5.** Plant Biomass of grassland communities by plant functional group under drought and ambient conditions in August of the last of 8 consecutive years of drought treatments (2016) and August one year after the last treatment (2017). Aboveground biomass was determined by cutting samples from a 0.1 $m^2$ subplot. The plant material was sorted into target species, weeds, and dead plant material, dried and weighed. A – biomass of grasses, B – biomass of legumes, C – biomass of herbs, D – results of ANOVAs to test for effects of the date, drought treatment and plant diversity on biomass of the plant functional groups.

**Supplementary Table S1.** Mean aboveground biomass of total plant biomass, grasses, legumes, and herbs under ambient (control) and drought conditions and results of paired t-test comparing biomass of drought to control.

| | **Mean biomass [g]** | | **t.test** | |
|---|---|---|---|---|
| | **Drought** | **Control** | **T** | **p** |
| **Total Plant Biomass** | 167.76 | 174.23 | -0.64 | 0.53 |
| **Grasses** | 27.46 | 28.50 | -0.21 | 0.83 |
| **Legumes** | 37.13 | 31.40 | 0.63 | 0.53 |
| **Herbs** | 56.93 | 62.48 | -0.89 | 0.37 |

**Supplementary Table S2.** ANOVA of drought effects and effects of plant community composition on soil water content (SWC) and aboveground plant biomass.

| | **SWC** | | | **Aboveground Plant Biomass** | | |
|---|---|---|---|---|---|---|
| | **Df** | **F value** | **P value** | **Df** | **F value** | **P value** |
| Drought Treatment | 1 | 9.55 | **0.00** | 1 | 0.41 | 0.53 |
| SWC | - | - | - | 1 | 20.72 | **<0.01** |
| Plot | 74 | 8.07 | **<0.01** | 74 | 3.63 | **<0.01** |
| Residuals | 74 | | | 73 | | |
| **Drought** | **Df** | **F value** | **P value** | **Df** | **F value** | **P value** |
| Plant Diversity$_{log}$ | 1 | 12.12 | **<0.01** | 1 | 19.41 | **<0.01** |
| Herbs$_{P/A}$ | 1 | 0.13 | 0.72 | 1 | 1.17 | 0.28 |
| Grasses$_{P/A}$ | 1 | 0.49 | 0.48 | 1 | 2.35 | 0.13 |
| Legumes$_{P/A}$ | 1 | 0.94 | 0.34 | 1 | 6.21 | **0.02** |
| Block | 3 | 8.73 | **<0.01** | 3 | 1.59 | 0.19 |
| Residuals | 67 | | | 67 | | |
| **Control** | **Df** | **F value** | **P value** | **Df** | **F value** | **P value** |
| Plant Diversity$_{log}$ | 1 | 22.49 | **<0.01** | 1 | 8.19 | **<0.01** |
| Herbs$_{P/A}$ | 1 | 2.93 | 0.09 | 1 | 2.17 | 0.14 |
| Grasses$_{P/A}$ | 1 | 2.42 | 0.12 | 1 | 0.69 | 0.41 |
| Legumes$_{P/A}$ | 1 | 1.49 | 0.23 | 1 | 5.98 | **0.02** |
| Block | 3 | 16.29 | **<0.01** | 3 | 0.93 | 0.43 |
| Residuals | 67 | | | 67 | | |

**Supplementary Table S3.** ANOVA of alpha-diversity (VT or ASV richness) in AMF and total fungi community as response to drought treatment and soil water content (SWC) stratified to plot, and as response to plant community composition stratified to block.

| | **AMF VT richness** | | | | **Total fungi ASV richness** | | | |
|---|---|---|---|---|---|---|---|---|
| | **Df** | **F value** | **$R^2$** | **P value** | **Df** | **F value** | **$R^2$** | **P value** |
| Drought Treatment | 1 | 0.25 | 0.00 | 0.60 | 1 | 1.39 | 0.01 | 0.12 |
| SWC | 1 | 3.20 | 0.02 | 0.29 | 1 | 0.09 | 0.00 | 0.39 |
| Residuals | 149 | | 0.99 | | 149 | | 0.99 | |
| **Drought** | **Df** | **F value** | **$R^2$** | **P value** | **Df** | **F value** | **$R^2$** | **P value** |
| Plant Diversity$_{log}$ | 1 | 0.52 | 0.01 | 0.46 | | 1.78 | 0.02 | 0.18 |
| Herbs$_{P/A}$ | 1 | 0.78 | 0.01 | 0.39 | 1 | 2.45 | 0.03 | 0.12 |
| Grasses$_{P/A}$ | 1 | 0.38 | 0.01 | 0.58 | 1 | 0.29 | 0.00 | 0.59 |
| Legumes$_{P/A}$ | 1 | 0.73 | 0.01 | 0.39 | 1 | 5.33 | 0.07 | **0.02** |
| Funct. Groups | 3 | 2.27 | 0.03 | 0.17 | 3 | 0.72 | 0.01 | 0.40 |
| Residuals | 69 | | 0.94 | | 69 | | 0.87 | |
| **Control** | **Df** | **F value** | **$R^2$** | **P value** | **Df** | **F value** | **$R^2$** | **P value** |
| Plant Diversity$_{log}$ | 1 | 1.48 | 0.02 | 0.22 | 1 | 1.79 | 0.02 | 0.18 |
| Herbs$_{P/A}$ | 1 | 5.08 | 0.07 | **0.02** | 1 | 0.94 | 0.01 | 0.32 |
| Grasses$_{P/A}$ | 1 | 0.42 | 0.01 | 0.54 | 1 | 0.81 | 0.01 | 0.35 |
| Legumes$_{P/A}$ | 1 | 0.18 | 0.00 | 0.72 | 1 | 3.34 | 0.04 | 0.08 |
| Funct. Groups | 3 | 0.06 | 0.00 | 0.89 | 3 | 0.47 | 0.01 | 0.50 |
| Residuals | 69 | | 0.90 | | 69 | | 0.90 | |
